# Synthetic reconstitution of planar polarity initiation reveals collective migration as a symmetry-breaking cue

**DOI:** 10.64898/2026.02.23.707595

**Authors:** Leah A Wallach, Connor D Thomas, Pulin Li

## Abstract

Planar cell polarity (PCP) organizes cellular behaviors along tissue axes, yet how symmetry is broken to initiate polarity across initially symmetric tissues remains unknown. We reconstituted tissue-scale polarity emergence *ex vivo* using an engineered epithelial system. A collective migration cue induces front-back enrichment of CELSR junctions, establishing aligned multicellular polarity that tracks migration direction and speed. A migration-onset wave consistently precedes the polarity-initiation wave, indicating that individual cells directly interpret the global cue rather than acquire polarity through intercellular relay. Strikingly, CELSR polarity arises without reciprocal VANGL-FZD interactions, identifying CELSR as the driver for planar polarization. These findings reveal how a global migratory cue can break symmetry and coordinate PCP across molecular, cellular, and tissue scales, providing a framework for engineering polarized tissues.

## INTRODUCTION

Planar cell polarity (PCP), the asymmetric organization of intercellular junctional complexes within the plane of a tissue, is essential for orienting developmental processes along body and organ axes (*1, 2*). Disruption of PCP causes severe developmental defects, including craniorachischisis, spina bifida, chronic lung disease, and hearing loss (*2–6*). Despite its fundamental importance, how nonpolarized tissues break symmetry to initiate planar polarity remains unknown.

A central unresolved question is the nature of the symmetry-breaking cue that initiates PCP. Polarity must be established at the single-cell level and aligned across millimeter-scale tissues, presenting a unique multiscale patterning problem (Fig. 1A) (*7*). Spatial concentration gradients of proteins associated with the PCP pathway, such as WNT and Frizzled (FZD) receptors, were leading candidates for polarity orienting signals, but are now disputed (*8–10*). Mechanical or force-based signals have been shown to maintain or re-orient the polarity axis in tissues that are already polarized, but whether these signals can induce junctional core PCP *de novo* remain unknown (*7, 10–16*).

**Figure 1.**
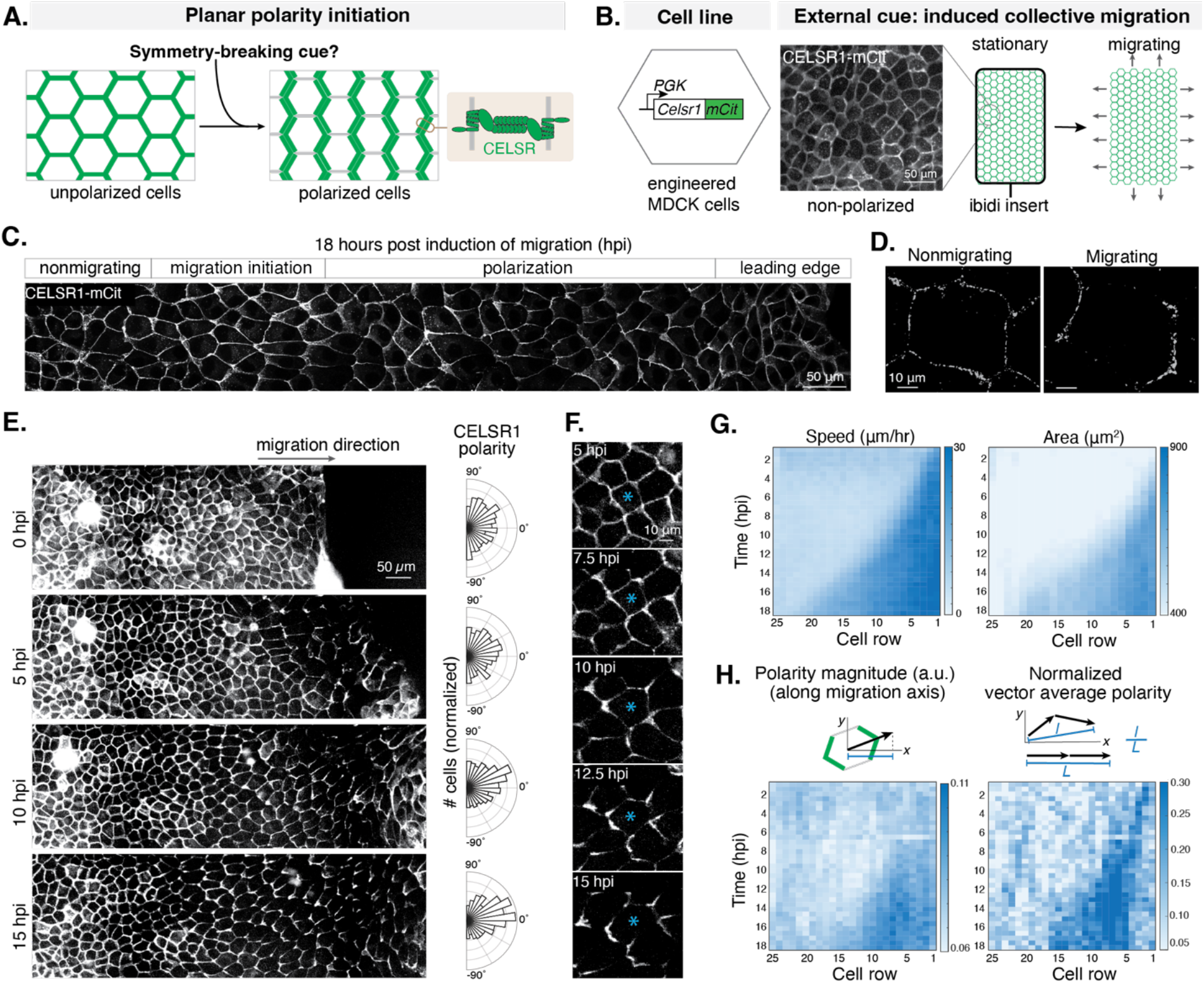
Collective migration breaks symmetry to induce planar polarity in engineered epithelial cells. **A**. Schematic of planar polarity initiation in an epithelial tissue induced by an unknown symmetry-breaking cue. **B**. Experimental setup for planar polarity reconstitution. *Left*: Engineered MDCK II clonal line overexpressing CELSR1-mCitrine showed uniform fluorescence distribution in stationary culture under confluency. *Right*: Induced collective migration assay, in which cells are grown to confluence in culture inserts and the removal of the insert triggers collective cell migration starting at the free edges of the field. **C**. Representative confocal image of CELSR1-mCitrine distribution after 18 hours post induction of migration (hpi). Maximum intensity projection of a 1.5 μm-thick Z stack. **D**. Distribution of CELSR1-mCitrine signal at the junctions of representative nonmigratory and migratory cells. Maximum intensity projection from 6 μm-thick Z stack. **E**. Epifluorescence imaging of a representative field of migrating CELSR1-mCitrine cells over time, with circular histograms (polar plots) showing orientation of CELSR1 polarity in migrating cells at the corresponding timepoints, scaled by polarity magnitude (1062 cells, n=3). **F**. Epifluorescence imaging centered on a single cell *(blue asterisk)*. **G-H**. Spatially and temporally binned kymographs showing dynamics of migration and polarity initiation in migrating MDCKs (25408 cell traces, n=3). CELSR1 polarity is measured by both polarity magnitude and normalized vector average polarity (defined as the ratio of the vector sum to the scalar sum of polarity vectors; see Methods). Both migration and polarity propagate as waves from the leading edge backward into the tissue.

Just as the identity of the symmetry-breaking cue remains unknown, its mode of action is also actively debated. In the “global cue” model, a tissue-wide signal acts simultaneously on all cells to initiate PCP, whereas the “local relay” model posits that polarity emerges in a small subset of cells and is subsequently propagated to neighboring cells through junctional interactions (*7, 17–20*). The intrinsic intercellular nature of PCP complexes, bridged across cell-cell junctions by homotypic interaction of the atypical cadherin CELSR, renders both models plausible (*2, 21, 22*).

However, distinguishing between them requires an experimental system in which symmetry breaking can be initiated in an otherwise unpolarized tissue and followed in real time, and in natural tissues, PCP emergence often coincides with massive tissue reorganization caused by parallel developmental events, complicating the mechanistic dissection of PCP emergence.

Here, we synthetically reconstituted tissue-scale planar polarity in initially nonpolarized epithelial cells and demonstrated that collective migration acts as a global symmetry-breaking cue to initiate and maintain planar polarity. Furthermore, CELSR can sense migration-induced cellular asymmetry to generate axial polarity independently of the feedback interactions between Frizzled (FZD) and Van Gogh (VANGL) across the junction. By connecting molecular dynamics to cellular asymmetry and tissue-scale patterning phenomena, these results provide a new framework for dissecting and engineering long-range multicellular polarity.

## RESULTS

### Collective cell migration induces tissue-scale symmetry breaking and planar polarity emergence

To isolate PCP from parallel developmental processes and test the sufficiency of external cues in breaking symmetry, we sought to reconstitute PCP in cultured cells. The intrinsically multicellular nature of PCP is best modeled by cells that readily form junctional cell-cell contacts. Therefore, we chose Madin-Darby Canine Kidney cells (MDCKs), a well-characterized model of epithelial tissues which naturally form tight and adherens junctions when cultured to confluency as a monolayer (*23, 24*). To directly observe polarity emergence through live imaging, we generated a stable clonal cell line that overexpresses mouse CELSR1 fused to mCitrine on the C-terminus, a fusion design that was previously validated using knock-in mice (*25*). Overexpressed CELSR1-mCitrine localized to cell-cell junctions, confirming the preservation of its natural homotypic interactions (Fig 1B).

Although CELSR1-mCitrine junctions appeared to be uniformly distributed along the cell boundaries without polarization in monolayer epithelia, when cells were grown to very high confluency, coordinated asymmetry of CELSR1 distribution emerged in localized regions (Fig S1). These regions had visible patterns of cell “swirling” based on their directionally elongated morphology, reminiscent of previously reported coordinated cell movement in confluent epithelial cells (*26*). Such coordinated movement, commonly referred to as tissue flow, was previously proposed to re-orient the polarity axis of tissues that are already planar polarized, such as the fly wing, raising an interesting possibility of coordinated movement acting as a symmetry-breaking cue to induce PCP initiation (*12, 13*).

To test this hypothesis, we forced cells to migrate in a classic collective migration assay. CELSR1-mCitrine+ cells were first cultured in a removable well insert to confluency and then released to induce linear collective migration (Fig 1B). By 18 hours post induction of migration (hpi), planar polarity of CELSR1 emerged along the axis of migration among migrating cells, excluding the leader cells (Fig 1C). While CELSR1 clusters were uniformly localized around the cell boundaries in nonmigrating cells, they were enriched to the front-rear faces and depleted from the left-right faces in migrating cells (Fig 1D). Importantly, CELSR1, in both unpolarized and polarized cells, was restricted to cell-cell contacts at lateral junctions, and was not visible either in basal structures or on apical cell surfaces (Fig S2, Supplementary Video 1). This pattern of CELSR1 is comparable to observed epithelial localization of PCP complexes *in vivo*, which similarly localize to cell-cell contacts rather than biased apical or basal protrusions (*2, 27–29*). Furthermore, neither endogenous CELSR1 nor low levels of CELSR1-mCitrine produced by a titratable system polarized during migration, indicating that a sufficiently high level of CELSR1 is necessary for migration-induced polarity (Fig S3B-D). Together, we demonstrate for the first time that collective migration can initiate a genuine asymmetric rearrangement of CELSR junctions, leading to the emergence of planar polarity from previously nonpolarized cells.

### Migration onset and polarity initiation propagate spatially as two travelling waves

The reconstituted system allowed us to quantify the spatiotemporal dynamics of migration and polarity emergence in individual cells and across the entire tissue simultaneously. Oriented CELSR1 asymmetry emerged and became coordinated over hundreds of cells, as measured by the polarity angle among all migrating cells (Fig 1E, F, Supplementary Video 2). By tracking and segmenting cells over 20 hrs of migration, we observed that CELSR1-mCitrine signal selectively intensified at the front-back junctions in individual cells once they started migration (Fig. 1F, Supplementary Video 3).

Furthermore, polarity emerged as a traveling wave across the tissue, closely following the onset of migration that propagated row by row. A wave of migration onset became apparent immediately upon release of the insert, originating from the leading edge and moving backwards into the tissue. This wave was manifested by coordinated increase in both migration speed and cell area (Fig. 1G). Polarity emerged subsequently as a second traveling wave, measured as an increase in both the polarity magnitude of individual cells and the alignment strength of cell polarity (Fig. 1H). Together, these results demonstrate that collective migration can induce large-scale, multicellular polarity in previously unpolarized tissues.

### The migration cue acts globally on every cell to break symmetry and initiate polarity

Although collective migration functions as a symmetry-breaking cue that propagates across hundreds of cells, the spatial scale at which such cues must operate within a tissue has long been debated, giving rise to the global-cue vs local-relay models (Fig 2A). While we observed that migration and polarity both propagate as two travelling waves side by side, this alone did not resolve whether migration serves as a global symmetry-breaking signal interpreted directly by each cell, or instead polarizes only the first few rows, with polarity subsequently propagating through an intercellular relay independently of continued tissue-scale movement. Discriminating between these mechanisms is essential for understanding whether symmetry breaking is continuously instructed by a tissue-wide cue, or becomes self-sustaining once locally triggered.

**Figure 2.**
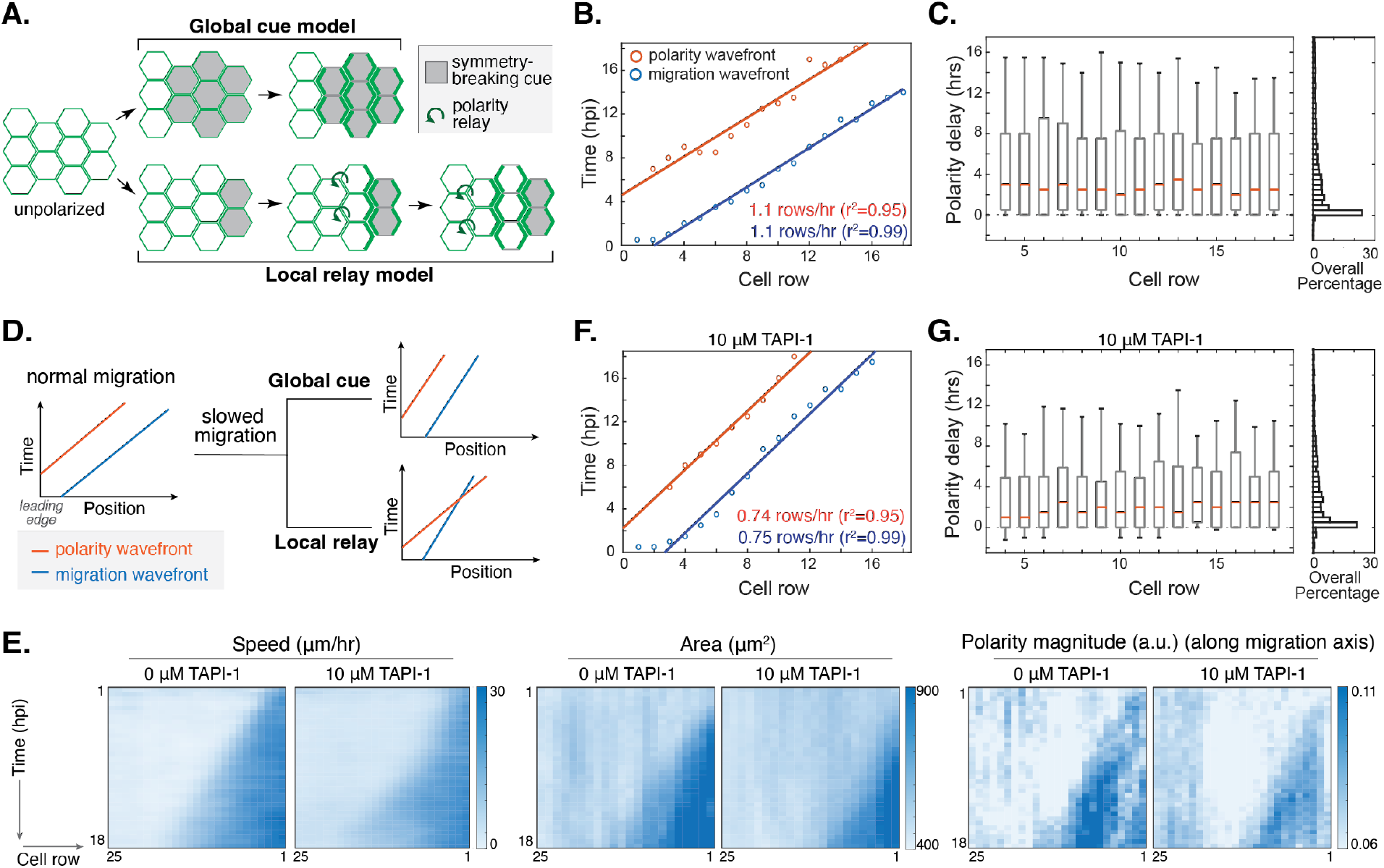
The migration cue acts globally on every cell to break symmetry and initiate polarity. **A**. Competing models describing the range of action of symmetry-breaking cues. **B-C**. Polarity initiation lags behind migration onset across the tissue. Markov models were applied to spatially binned (**B**) and single cell (**C**) data to determine initiation times for migration and polarization. **B**. The two waves travel at the same constant speeds across the tissue. Lines are linear fits of the wavefront positions (25408 cell traces, n=3). **C**. Polarity initiation has a mean time delay of 3.1 hrs behind migration onset in individual cells and the delay is position-invariant (256 cells per row, One-way ANOVA, p = 0.68). Leading rows were omitted due to the lack of polarity. Box plots indicate median and the 10, 25, 75 and 90 percentile marks. **D**. The two models predict different relationships between the two travelling waves when migration is slowed down. **E**. Kymographs of cell migration speed, cell area, and polarity magnitude from representative control (+0 µM TAPI-1) and slowed (+10 µM TAPI-1) migrating cell fields. **F & G**. Polarity initiation maintains a constant delay behind migration onset when migration slows down, using spatially binned data **(F)** (9276 cell traces, n=3) and individual cells **(G)** (170 cells per cell row), with a mean delay of 3.7 hrs that is position-invariant (One-way ANOVA, p = 0.93).

To distinguish between these models, we quantified the spatiotemporal relationship between migration onset and polarity initiation by live imaging. If migration acts as a global symmetry-breaking cue that each cell must experience prior to polarization, the migration and polarity waves should propagate into the tissue at the same speed with a fixed time delay. In contrast, a local-relay mechanism would permit different wave speeds and spatially variable delays. We spatially binned cells into rows and trained Markov models on migration and polarity metrics to infer the timing of migration onset and polarity initiation for each bin (Fig S4A-D). We found that migration onset and polarity initiation propagated at identical speeds, with polarity initiation consistently lagging migration (Fig 2B). Applying the same model at single-cell resolution confirmed this relationship: polarity initiation followed migration onset by an average delay of 3.1 hours, independent of a cell’s position within the tissue, consistent with the direct coupling of a global cue (Figs 2C, S4E).

To directly test if polarity initiation is causally coupled to migration onset in individual cells, we slowed migration using TAPI-1, an ADAM17 and matrix metalloprotease inhibitor (*27*). Migration speed decreased to 67% of untreated control (Fig S5A). Under the global-cue model, this perturbation should proportionally slow down the polarity wave, whereas the local-cue model predicts an unaltered polarity wave (Fig 2D). (*30*), Strikingly, CELSR1 polarization still propagated across as a traveling wave, but the polarity wave slowed in parallel with migration, with both waves moving at the same speed and maintaining a constant delay of 3.7 hours (Figs 2E-G, S5B-C). Thus, polarity initiation is directly coupled to migration onset in individual cells: each cell must migrate to break symmetry, rather than acquiring polarity from polarized neighbors. Because collective migration of epithelial sheets is inherently coordinated through adhesion and cytoskeletal coupling (*31*), the resulting traveling wave of migration synchronizes both the timing and axis of planar polarity initiation across the tissue.

### Sustained CELSR polarity requires an ongoing migration cue

Our discovery of migration-induced planar polarity emergence and the well-known function of PCP in coordinating large-scale cell movement suggest that migration and planar polarity may be dynamically coupled in certain tissue contexts (*32*). We therefore asked whether, once established, CELSR polarity becomes self-sustaining or instead remains dependent on ongoing migration. To address this question, we first asked if established CELSR polarity requires ongoing migration to maintain. We allowed cells to migrate for 16 hours and then acutely halted migration with an Arp2/3 inhibitor (400µM CK666), which synchronously stopped forward motion across the epithelial sheet (Fig 3A). Halted cells maintained their stretched morphology and junctional contact with neighboring cells, with CELSR1 remaining at cell-cell junctions (Fig 3C, S6A, Supplementary Video 4). Stalled cells resumed migration after inhibitor washout, demonstrating that drug treatment did not lastingly damage cell functions (Fig S6B). Strikingly, migration arrest led to rapid loss of both polarity strength and alignment, demonstrating that migration serves not only as an initiation cue but also a maintenance signal (Figs 3B, S6A).

**Figure 3.**
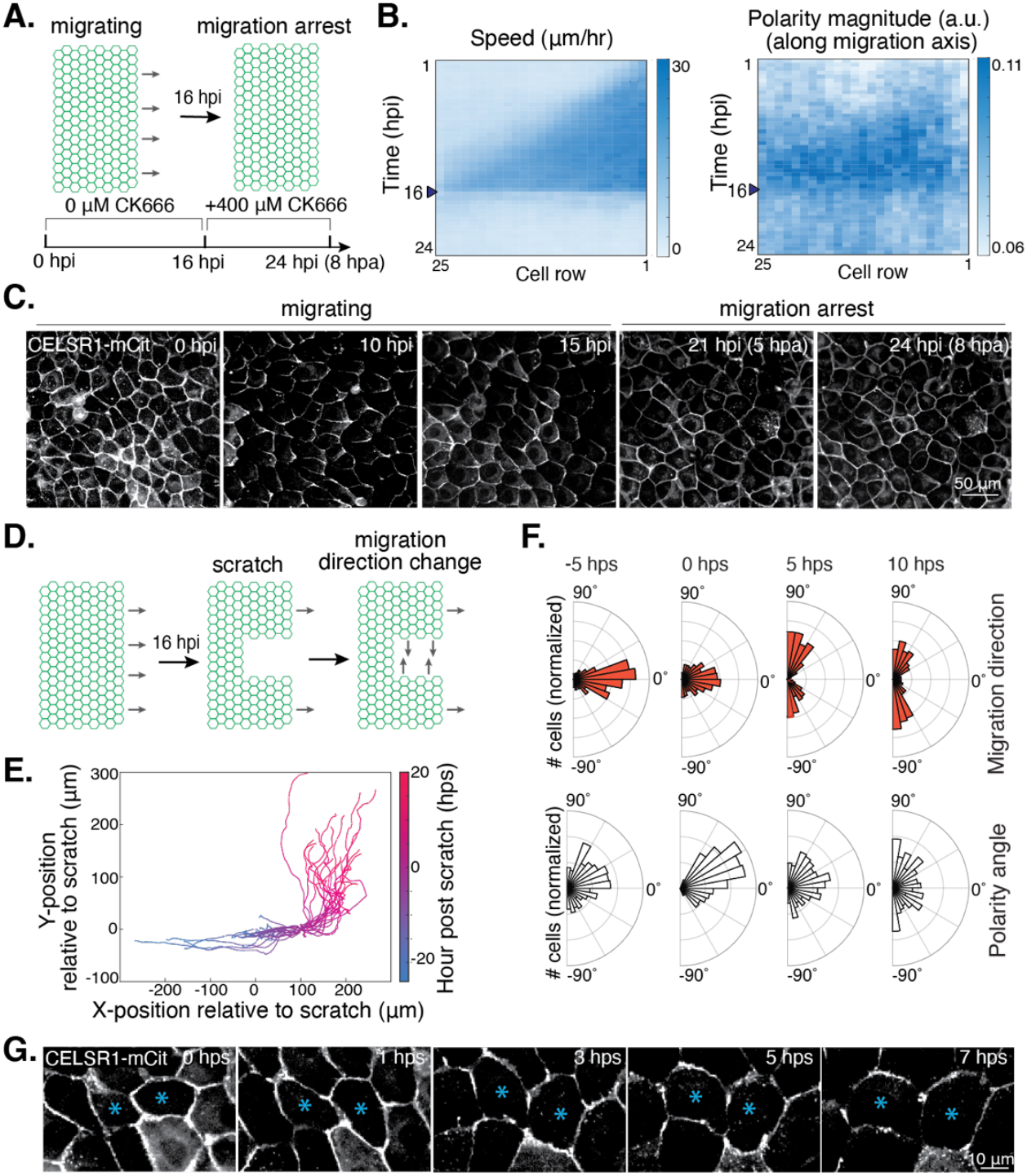
Sustained CELSR polarity requires an ongoing migration cue. **A**. Schematic of migration arrest experiments. Cells are allowed to migrate for 16 hrs before the addition of CK-666, an Arp2/3 inhibitor. hpi, hours post initiation; hpa, hours post arrest. **B**. Kymographs of migration speed and polarity. Within 2 hrs of CK-666 treatment, migration is fully arrested and CELSR1 polarity is largely lost (21246 cell traces, n=4). **C**. Timelapse images of a representative cell field show the loss of polarity upon migration arrest. **D**. Schematic of the migration reorientation experiment. Migrating cell fields were scratched at 24 hpi to introduce new free edges orthogonal to the initial migration direction. **E**. Migration trajectories show switching of directions. Each trace represents a single cell, with X and Y coordinates normalized to its position at 0 hour post scratch (hps). **F**. Polar plots of migration direction and polarity orientation after scratch. Only polarized migrating cells at 0 hps were included (396,459, 270 and 179 cells for respective time points, n=3). **G**. Timelapse images of a representative cell field show redistribution of CELSR1 protein in response to change in migration direction.

Next, we asked whether established polarity remains sensitive to the direction of ongoing migration by performing a reorientation assay (Fig 3D). In this assay, an already-migrating cell field was scratched in the orthogonal direction of the original open boundary, causing nearby cells to rapidly reorient and migrate perpendicularly to their previous migration direction (Figs 3E-F, S7A, Supplementary Video 5). Although cells further enlarged during re-orientation, they exhibited minimal neighbor exchange (Fig S7B, Supplementary Video 6). Remarkably, cells with pre-established polarity reoriented their polarity along the new migration direction within 5 hours, at both single-cell and tissue scales (Fig 3F, G, Supplementary Video 6). Together, these results show that CELSR continuously reads out the presence and direction of collective migration to maintain or adjust its polarity, consistent with previously observed migration-induced reorientation in the *Drosophila* wing (*12, 13*).

### Polarity is not a general feature of cell junctions

Although overexpressed CELSR polarizes across the tissue, it is important to determine whether this reflects a specific property of CELSR or a general feature of membrane or junctional proteins responding to migration. To address this, we compared CELSR behavior to that of other membrane-associated proteins. In cells coexpressing CELSR1-mScarletI and a generic membrane-associated protein (mCitrine-CAAX), only CELSR1-mScarletI polarized along the migration axis, whereas mCitrine-CAAX remained uniformly distributed (Fig 4A). This result confirms that polarity is not an artefact of either high-level mCitrine expression or bulk membrane flow.

**Figure 4.**
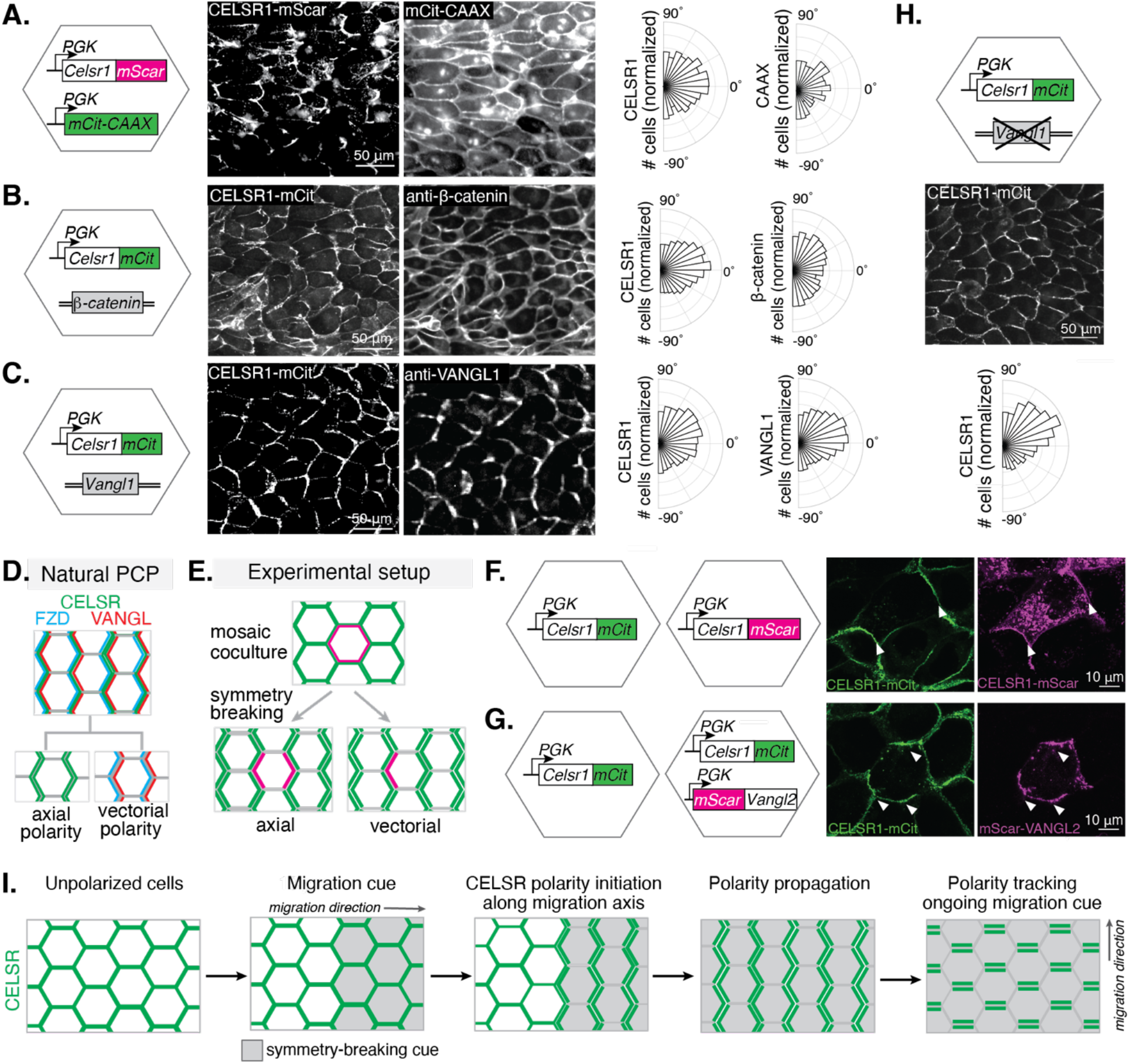
Polarization is CELSR-dependent and CELSR-specific. **A**. Polarity comparison of CELSR1-mScarletI and mCitrine-CAAX (generic membrane-associated protein) co-expressed in the same cell at 18 hpi (2247 cells, n=3). **B**. Polarity comparison of CELSR1-mCitrine and endogenously expressed β-catenin (anti-β-catenin immunostaining) at 18 hpi (5358 cells, n=3). **C**. Polarity comparison of CELSR1-mCitrine and endogenously expressed VANGL1 (anti-VANGL1 immunostaining) protein in the same cells at 18 hpi (5235 cells, n=2). **D**. Schematic of the classic PCP pattern. CELSR presents at both front and rear sides of polarized cells, while VANGL and FZD localize to opposing sides, defining a vector of polarity along the CELSR polarity axis. **E-G**. Mosaic coculture analysis to distinguish between axial and vectorial polarity. Individual cells expressing mScarletI-tagged CELSR1 or VANGL2 are surrounded by cells expressing only CELSR1-mCitrine (**E**). Representative images of a polarized CELSR1-mScarletI cell (**F**) and a polarized mScarletI-VANGL2 cell (**G**), with mScarletI detected on both the front and rear sides of the cell, consistent with axial polarity. White arrowheads denote regions of polarized protein localization. **H**. Polarization of CELSR1-mCitrine in *Vangl1* KO MDCK cells at 18 hpi (1208 cells, n=2). **I**. Model of CELSR planar polarity initiation.

We next asked whether other junctional proteins participate in migration-induced polarity. We found that endogenous β-catenin, a core component of adherens junctions, did not polarize along the migration axis in CELSR1-mCitrine+ cells, consistent with *in vivo* observations that PCP proteins polarize independently of adherens junctions (Fig 4B) (*27*). Moreover, overexpressed E-cadherin-mScarletI failed to polarize in response to migration, confirming that the lack of polarization was not due to insufficient protein levels (Fig S8). Together, these results demonstrate that migration-induced polarization is not a general property of cell junctions but is specific to core PCP, suggesting that collective migration acts as a PCP-selective symmetry-breaking cue.

### CELSR1 drives axial polarity independently of vectorial polarity

We next asked if migration-induced CELSR1 polarization requires the complete set of PCP core components. In natural tissues, PCP establishes both axial polarity (alignment along a tissue axis) and vectorial polarity (asymmetric segregation of components such as FZD and VANGL to opposite cell edges) (Fig 4C) (*2, 7*). Molecular feedback interactions between FZD and VANGL are at least partly responsible for vectorial polarity (*1, 2, 33–35*). Whether the emergence of CELSR1 axial polarity depends on the concurrent establishment of vectorial polarity remains unclear.

In polarized CELSR1-Citrine overexpression cells, both endogenous VANGL1 and overexpressed mScarletI-VANGL2 were also polarized during collective migration (Figs 4C, S9A). This is in stark contrast to endogenously expressed FZD6 and VANGL1, neither of which polarized in migrating wild-type cells; neither did overexpressed mScarletI-VANGL2 alone (Figs S3A, S9B, Table S1). These results suggest that CELSR is required to drive polarization, but VANGL is not, even at high expression levels.

To ask whether the polarity of CELSR and VANGL is axial only, or both axial and vectorial, we established a mosaic co-culture assay that allows us to distinguish the junctional proteins contributed by each of the neighboring cells (Fig 4D). Mixing CELSR1-mCitrine MDCKs and sparse CELSR1-mScarletI MDCKs, we isolated individual mScarletI+ cell and confirmed the axial polarity of CELSR1-mScarletI (Fig 4E). Similarly, diluting CELSR1-mCitrine/mScarletI-VANGL2 cells with CELSR1-mCitrine cells revealed VANGL2 localization at both the front and back edges of individual polarized cells, consistent with axial rather than vectorial polarity (Fig 4F). Importantly, CELSR1-mCitrine cells retained highly coordinated tissue polarity in response to migration after complete knockout of endogenous *Vangl1* (Figs 4G, S9C-D). These findings demonstrate that migration-induced axial polarity emerges independently of vectorial PCP feedback. Thus, CELSR can directly interpret migratory cues to establish tissue-scale axial polarity without requiring reciprocal asymmetric segregation of FZD and VANGL.

## DISCUSSION

Planar cell polarity (PCP) provides a conserved mechanism for orienting cells and tissues during development. Decades of work across model organisms have established a detailed molecular framework describing how PCP is maintained and aligned once established (*1, 2, 36, 37*). In contrast, how initial symmetry is broken and how that event is coordinated across the tissue along a single axis has remained a fundamental unresolved question. Using a synthetically reconstituted epithelial tissue system, we demonstrate that induced collective migration is sufficient to break CELSR1 symmetry and initiate planar polarity across the tissue, underscoring the value of bottom-up reconstitution for uncovering minimal rules governing multicellular self-organization.

Our results support a new model of PCP initiation in which collective migration acts as a global symmetry-breaking cue (Fig 4H). Individual cells sense the migratory cue and redistribute CELSR to the front and back edges relative to the direction of migration, thereby establishing axial polarity. Migration then acts as a continuous instructive input that both orientates and maintains this polarity. How migration breaks symmetry within each cell, and how CELSR interprets this asymmetry into planar polarity, require future investigation. Furthermore, collective migration is unlikely to be the sole PCP-initiating cue. Across tissue contexts, PCP can respond to diverse mechanical perturbations, including tissue stretch and boundary-imposed forces, and not all planar polarized tissues are known to exhibit early migratory behavior (*2, 7, 10, 14, 16, 38, 39*). Therefore, we anticipate that other types of mechanical cues might similarly induce PCP initiation. Nevertheless, our findings provide a general mechanism by which tissue-scale mechanical asymmetries can be interpreted at the single-cell level to generate coordinated polarity.

The requirement for ongoing migration to sustain CELSR polarity further suggests that planar polarity is dynamically coupled to active morphogenetic processes. PCP frequently functions during transient morphogenic events, such as neural tube closure, kidney tubule elongation, and vertebrate limb bud elongation (*32, 40, 41*). Together with the established role of PCP in orienting cell migration, our findings support a model in which developmental morphogenetic events involve feedback between PCP and the orientation of morphogenetic forces, rather than a unidirectional signaling event (*42–44*). This raises the possibility that cessation of morphogenic events may destabilize PCP, and that additional stabilizing mechanisms may be required in tissues where PCP persists after the morphogenic event has ceased.

Finally, our results identify CELSR as a central mediator of cellular symmetry-breaking in PCP, and demonstrate that axial polarity can arise independently of vectorial polarity. This challenges prevailing models in which junctional asymmetry of PCP components such as FZD or VANGL is required for polarity emergence (*7, 18, 19, 45, 46*). One possibility is that CELSR-mediated axial polarity acts upstream of junctional asymmetry, concentrating other PCP components and facilitating subsequent *cis* and *trans* feedback interactions. Alternatively, junctional asymmetry, and thus vectorial polarity, may arise independently or concurrently through additional cues that are not present in our system. Notably, although CELSR is broadly required for mammalian PCP, partial polarity can arise independently of CELSR in *Drosophila*, suggesting evolutionary flexibility in symmetry-breaking mechanisms (*47, 48*). Extending synthetic reconstitution approaches to induce FZD–VANGL junctional asymmetry in engineered tissues will be essential for defining how planar symmetry-breaking is coupled to the canonical PCP junctional organization.

## Acknowledgements

We would like to thank Adam Martin, Peter Reddien, and Daniel Lew for valuable advice and support, in addition to members of the Li Lab for discussion and feedback through all stages of the project. We would like to thank Danielle Devenport for providing critical reagents. We would also like to thank the flow cytometry core facilities (Whitehead Institute and Koch Institute) and the W.M. Keck Microscopy Facility (Whitehead Institute) for providing access to their instruments, and the genome technology core (Whitehead Institute) and bioinformatics core (Whitehead Institute) for assistance in collecting RNA sequencing data. This work was supported by National Institute of Health grants DP2HD108777 (P.L.), Allen Family Philanthropies (P.L.), Eugene Bell Career Development Professorship (P.L.), the National Science Foundation Graduate Research Fellowship (L.W.). C.T. is a Valhalla Graduate Student Scholar.

## Competing interests

Authors declare that they have no competing interests.

## Author contributions

L.W. and P.L. conceived the project and designed the experiments. L.W. and C.T. performed the experiments and analyzed the data. L.W., C.T., and P.L. wrote the paper.

## Materials and Methods

### Constructs

Celsr1-mCitrine was cloned from the PEGFPN1-Celsr1-GFP construct generously provided by Danelle Devenport (*27*) through restriction digestion with HindIII and NheI, and inserted into a PiggyBac transposase backbone containing an MCS followed by a C-terminal mCitrine. Additional Celsr1 constructs were generated by HindIII/XbaI restriction digest of Celsr1-mCitrine to replace the C-terminal Citrine with mScarletI (cloned from 3xnls-mScarletI, a gift from Dorus Gadella, Addgene #98816) (*49*) or with Citrine-DHFR (*50*). PiggyBac-mScarletI-Vangl2 was cloned from mCherry-Vangl2 (generously provided by Danelle Devenport) (*27*). PiggyBac-E-cadherin-mScarletI was cloned from mouse E-Cadherin-GFP (a gift from Alpha Yap, Addgene #67937) (*51*). All plasmids generated were verified with Plasmidsaurus Nanopore sequencing.

### Cell line generation

MDCK IIs, a gift from the Jianzhu Chen lab, were cultured in MEM media (MEM with 10% Fetal Bovine Serum (FBS), 1% pen-strep-glutamine) and passaged every 3 days at 1:10 dilution. Briefly, cell lines were generated through PiggyBac transposase co-transfection with PiggyBac vector constructs using Lipofectamine LTX (*52*). After 24 hours of transfection, cells were selected with antibiotics for stable integration and sorted as single cells by FACS. Clones were chosen based on successful confluent monolayer formation and regular morphology comparable to wild-type cells. For all assays, cells were grown to confluence in wells or Ibidi 2-well culture inserts before imaging or insert release to induce migration.

### Immunofluorescence

Cultured cells were fixed in 4% PFA, permeabilized with PBSTx (PBS with 0.5% Triton-X), and blocked in NGS-PBST (PBS, 0.1% Tween20, 10% Normal Goat Serum) for 1.5 hours at room temperature. After overnight 4°C primary antibody incubation, samples were washed three times in PBST and incubated in secondary antibody, DAPI, and/or Phalloidin-594 at 25°C with rocking for one hour. Samples were washed three times in PBST and once in PBS, and imaged in PBS.

Primary antibodies used were: rabbit anti-Celsr1 (cat# SAB4503681, 1:200 dilution); rabbit anti-Vangl1 (cat# HPA025235, 1:200 dilution); goat anti-Fzd6 (cat# AF1526, 1:100 dilution); mouse anti-beta catenin (cat# 2677, 1:400 dilution). All primary antibodies were diluted in NGS-PBST blocking buffer. Secondary antibodies were diluted 1:4000 in blocking buffer.

### Image acquisition

For imaging, MDCK cells were cultured on glass- or glass-like plastic-bottomed 24-well imaging plates. Live cells were cultured in imaging media to reduce background fluorescence (DMEM FluoroBrite with 10% FBS, 1% pen-strep-glutamine). Fixed cells were imaged in PBS. Tissue-scale live and fixed images were collected on a Nikon Ti2 inverted epifluorescence microscope controlled by NIS-Elements using a 20x Plan Fluor NA. 5 objective. High-spatial-resolution images of Celsr1 distribution and other cellular markers were collected on a Zeiss LSM 980 confocal microscope with Airyscan controlled by Zen Blue 3.5. A Zeiss Plan Apo 63x/1.4 oil-immersion objective was used.

### Image analysis

Cells were segmented and tracked using the Cellpose-Trackmate plug-in in FIJI/ImageJ (*53, 54*). Cells were segmented with the Cytoplasm 2 pretrained model and tracked using a linear Kalman model. Segmentation masks were generated using FIJI/ImageJ. The resulting masks were analyzed with QuantifyPolarity to measure polarity statistics (*55*). Follow-up analysis was performed using custom Matlab scripts. Polar plots were generated with the Matlab polarhistogram function, using the ‘probability’ scaling function to normalize cell counts between plots. Oriented polarity angle was calculated by taking the cosine of the angle between the polarity direction and the migration direction of the cell field. Polarity magnitude along the migration axis was calculated by multiplying the polarity magnitude by the oriented polarity angle. Normalized vector average polarity was calculated by dividing the vector sum of selected cells by the sum of their magnitudes, resulting in a score between 0 and 1 that corresponds to directional coordination between vectors, as previously described (*55*). Cell binning was accomplished by dividing the migrating cell field into 25 bins of equal cell number of rank order distance to the leading edge. Pre-migration, these bins roughly equate to 1.5 cell widths.

### Determining timing of polarity and migration initiation

To determine the timing of migration onset and polarity initiation, we established a consistent approach to draw cutoffs between the respective states (nonmigratory vs migratory, nonpolarized vs polarized). In brief, we reasoned that stationary cells represented a true unpolarized population and used their polarity metrics to set cutoffs for binarizing the time series of polarity metrics among migrating cells. These binarized traces were then used to train a Markov model to infer whether observed polarity metrics are reflective of a polarized state and to predict the timing of state transition for both binned and single cell data.

In order to identify nonmigrating cells in binned data, we fit the distribution of mean X-displacement (defined as the distance traveled along the migration axis between the current and previous frame) to two Gaussian distributions. This process identified a distribution centered at zero displacement, corresponding to no net movement, which was assigned as the nonmigratory population. Bins with x-displacement less than two standard deviations above the mean of the nonmigratory population were labeled as stationary and used as baselines for expected nonpolarized behavior. The overall polarity magnitude, polarity magnitude along the migration axis, and normalized vector average polarity for stationary bins were well-described by a single Gaussian distribution. Polarity thresholds were set as the mean polarity values within the stationary population plus 1.5 standard deviations.

Binarized data for x-displacement and each of the three polarity metrics were used in conjunction with built-in Matlab hidden Markov modeling to train a two-state Markov model (Fig. S4A). A trace was started in an “off” state, corresponding to an unpolarized or nonmigratory cell state. At each timestep, it could either remain in this state or transition into an “on” state corresponding to having initiated migration or polarization. Data collected from each timestep is considered an emission, biased towards being above or below the thresholds based on underlying state. Resulting trained rates were used to decode each bin’s most likely state sequence based on their emission sequence and to identify timepoints at which the track switched between states. To ensure genuine polarity, bins were predicted to be polarized only upon consensus of at least two of the three models trained on polarity magnitude, polarity magnitude along the migration axis, and normalized vector average polarity. The first frame labeled as polarized for a given row was labeled as the initiation timepoint for that row. To achieve single-cell resolution, models trained using the polarity magnitude and speed values from binned data were applied to single-cell traces, identifying the frame of initiation for polarization and migration respectively.

To identify reorienting cells in the directional-change scratch assay, only cells within 300 μm of the scratch location and more than 100 μm from the original leading edge were considered. Cells were additionally required to be migrating (speed > 2 STD above pre-migration population, as in previous analysis) and polarized (polarity magnitude along initial migration axis > 0.1) one frame before the scratch, to limit analysis to polarity changes and not include new polarity initiation events.

### Generation of *Vangl1* knockout cell line

CELSR1-mCitrine MDCKs were CRISPR edited using the Alt-R™ CRISPR-Cas9 system (Integrated DNA Technologies). gRNA (CCCCATCAGGATGACAACTG) targeted Exon 4 of the endogenous *Canis familiaris VANGL1* (Chromosome 17, NC_051821.1 54055046..54108648). Alt-R™ crRNA containing the guide sequence was complexed with ATTO 647 tracrRNA and Alt-R Cas9 protein using manufacturer protocols. Successfully transfected cells (ATTO 647-positive) were FACS sorted as individual cells at 48 hours after transfection and grown for two weeks. Cells with proper morphology and continued CELSR1-mCitrine expression were selected and genotyped by PCR amplification of the targeted locus, using PCR primers TCTAGGGTTGCTCATGGGTG (forward) and ATGCGAACCCCGTAGAAGAG (reverse). Oxford Nanopore sequencing (Plasmidsaurus) identified edited alleles and confirmed complete knockout of both alleles. Loss of VANGL1 expression was confirmed by staining with VANGL1 antibody (cat# HPA025235, 1:200) and imaged on Nikon Ti2.

### RNA sequencing

Wild-type MDCK II cells were grown to confluence and collected for total mRNA purification. Library was constructed according to standard Illumina protocols and sequenced on NovaSeq 6000.

### Resource availability

#### Materials availability

plasmids and cell lines generated in this study will be available upon request. Data and code availability: Analysis codes are available at https://github.com/Pulin-Li-Lab/Planar_Cell_Polarity_Analysis

#### Lead contact

Requests for further information and reagents should be directed to the lead contact, Pulin Li, at.

---

**Figure S1.**
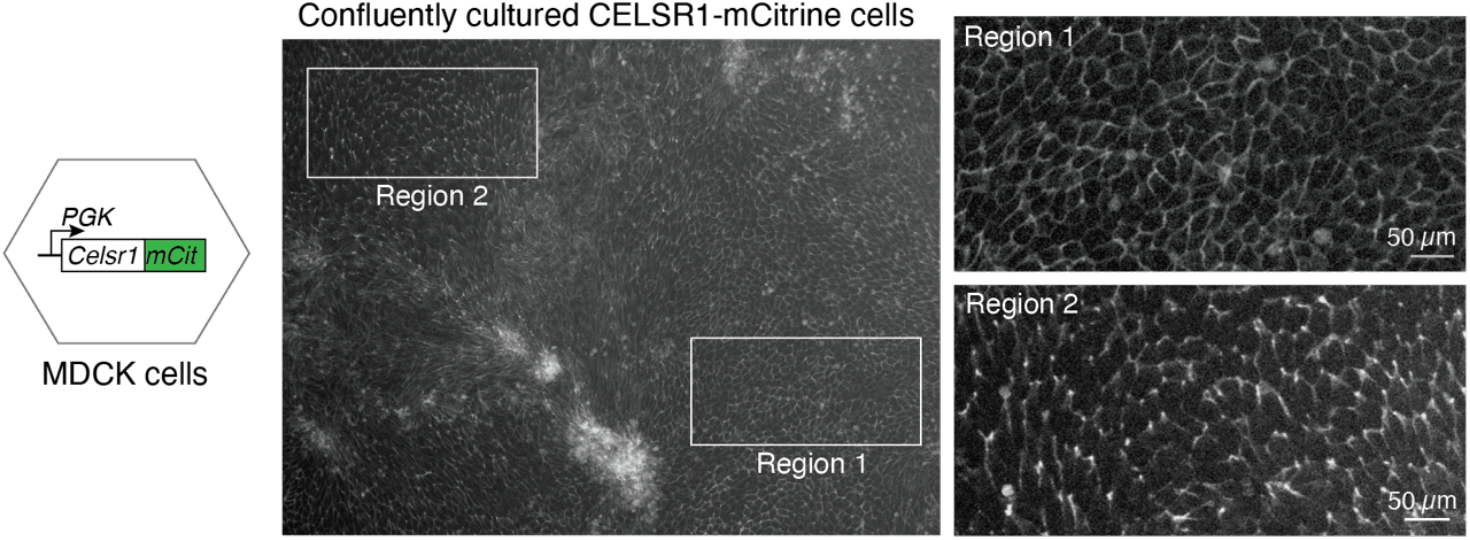
Emergence of CELSR1 planar polarity in engineered MDCK at high confluency. A clonal population of CELSR1-mCitrine MDCK cells were grown for 8 days after reaching confluency. While most regions of the well showed uniform distribution of CELSR1-mCitrine signal (Region 1), selective regions displayed organized asymmetric localization of mCitrine signal to certain cell junctions, corresponding to a planar polarized appearance (Region 2). Regions of cells with elongated morphology (Region 2) indicate tissue flow or collective migration, which is often observed in high-density cell culture.

**Figure S2.**
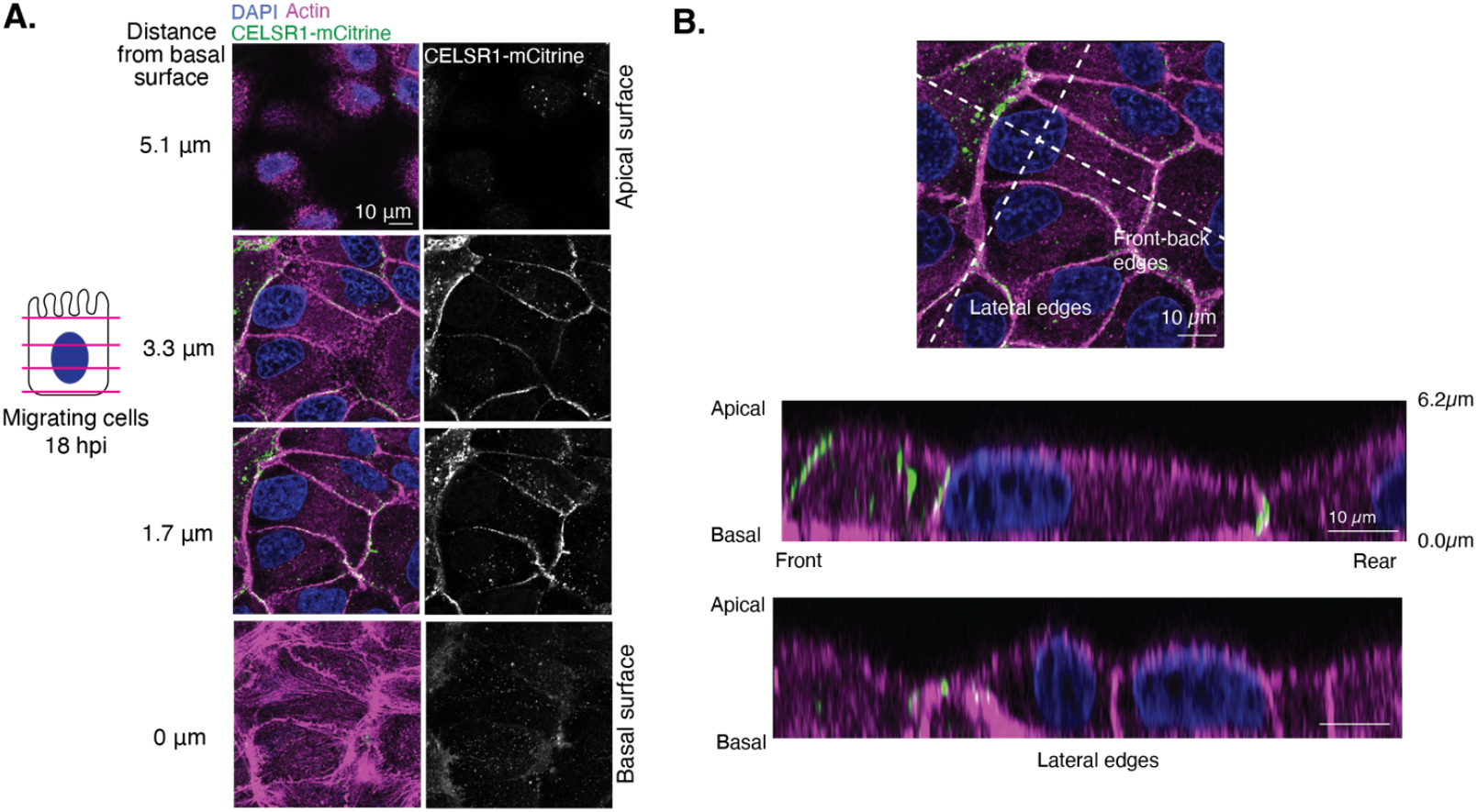
Characterization of CELSR1 subcellular localization in polarized migrating cells. **A**. Migrating CELSR1-mCitrine+ MDCK cells stained for nuclei (DAPI) and actin (Phalloidin-549) for cell boundaries. Confocal Z-slicing through polarized cells showed that CELSR1 is largely restricted to the lateral sides of cells corresponding to cell-cell contacts, with little to no CELSR1-mCitrine signal at the basal or apical cell surfaces. **B**. Orthogonal views through the front-rear and lateral sides of migrating cells show CELSR1 is polarized through preferential localization to front-back cell-cell junctions, but is not present in polarized basal or apical structures.

**Figure S3.**
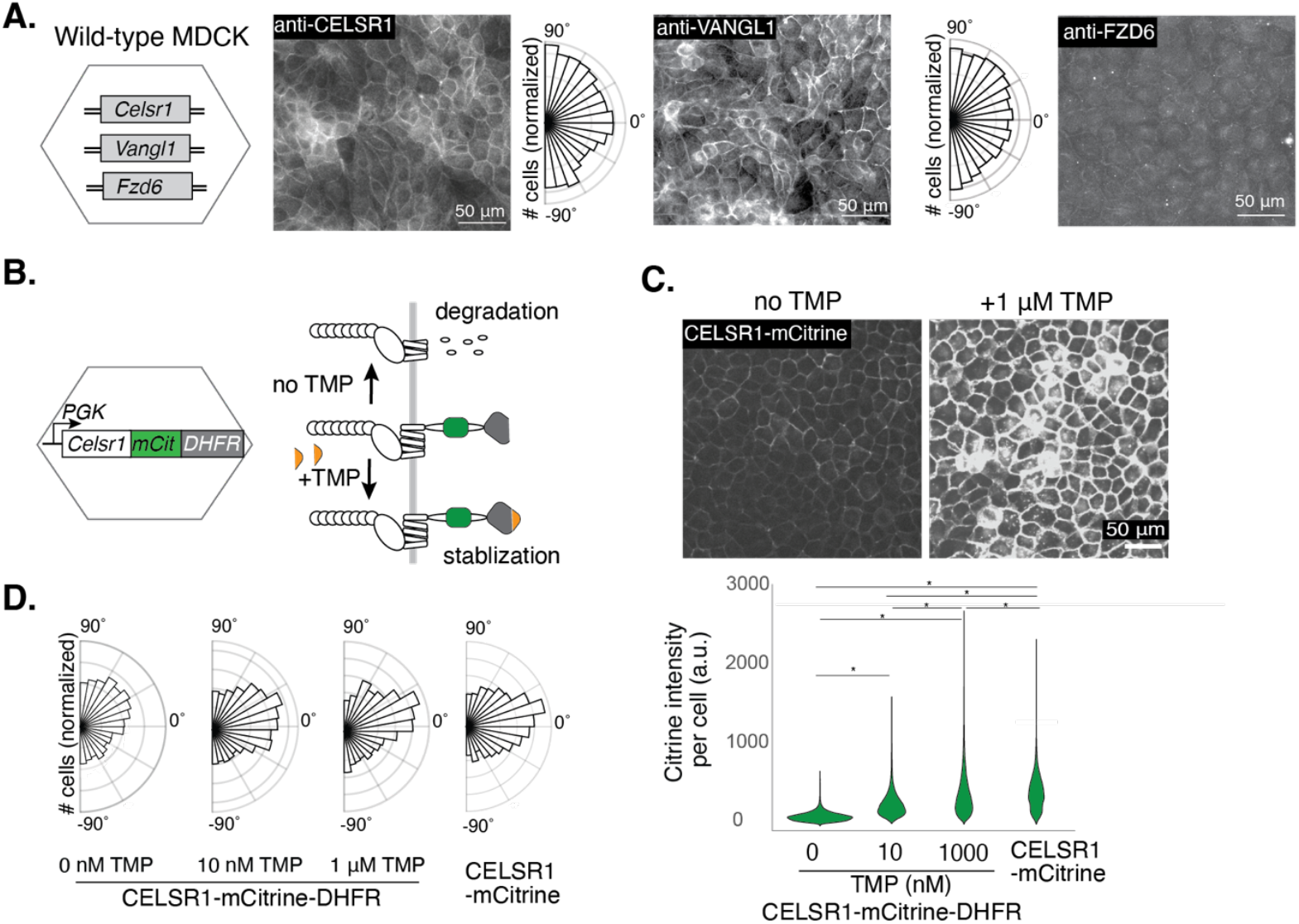
Polarization requires high levels of CELSR1 expression. **A**. Immunofluorescence staining for endogenous CELSR1, VANGL1, and FZD6 protein in migrating untransfected MDCK cells. Circular histograms show endogenous CELSR1 and Vangl1 polarity. Due to low signal, FZD6 could not be segmented or quantified for polarized distribution. **B**. Schematic of dose-titrable CELSR1-mCitrine-DHFR cell line. The intrinsically destabilized DHFR domain led to the degradation of the majority of CELSR1-mCitrine proteins. Addition of DHFR-binding drug trimethoprim (TMP) stabilized DHFR in a dose-dependent manner, leading to titratable CELSR1-mCitrine protein levels. **C**. Increasing doses of TMP led to increased CELSR1-mCitrine fluorescence signal. *Top*, representative images of untreated or TMP-treated cells. *Bottom*, quantification of Citrine intensity under different TMP doses, compared to the cell line that constitutively produces CELSR1-mCitrine without DHFR tag. * denotes statistical significance, p < 0.05 (One-way Anova and Tukey’s test). **D**. Increasing doses of TMP increased CELSR1 polarization during migration, with polar plots showing orientation of CELSR1 polarity in migrating cells at 18 hpi, scaled by polarity magnitude. (0 nM TMP, 5410 cells, 3 replicates; 10 nM TMP, 4937 cells, 3 replicates; 1 μM TMP, 3768 cells, 2 replicates; CELSR1-mCitrine, 25408 cell traces, 3 replicates)

**Figure S4.**
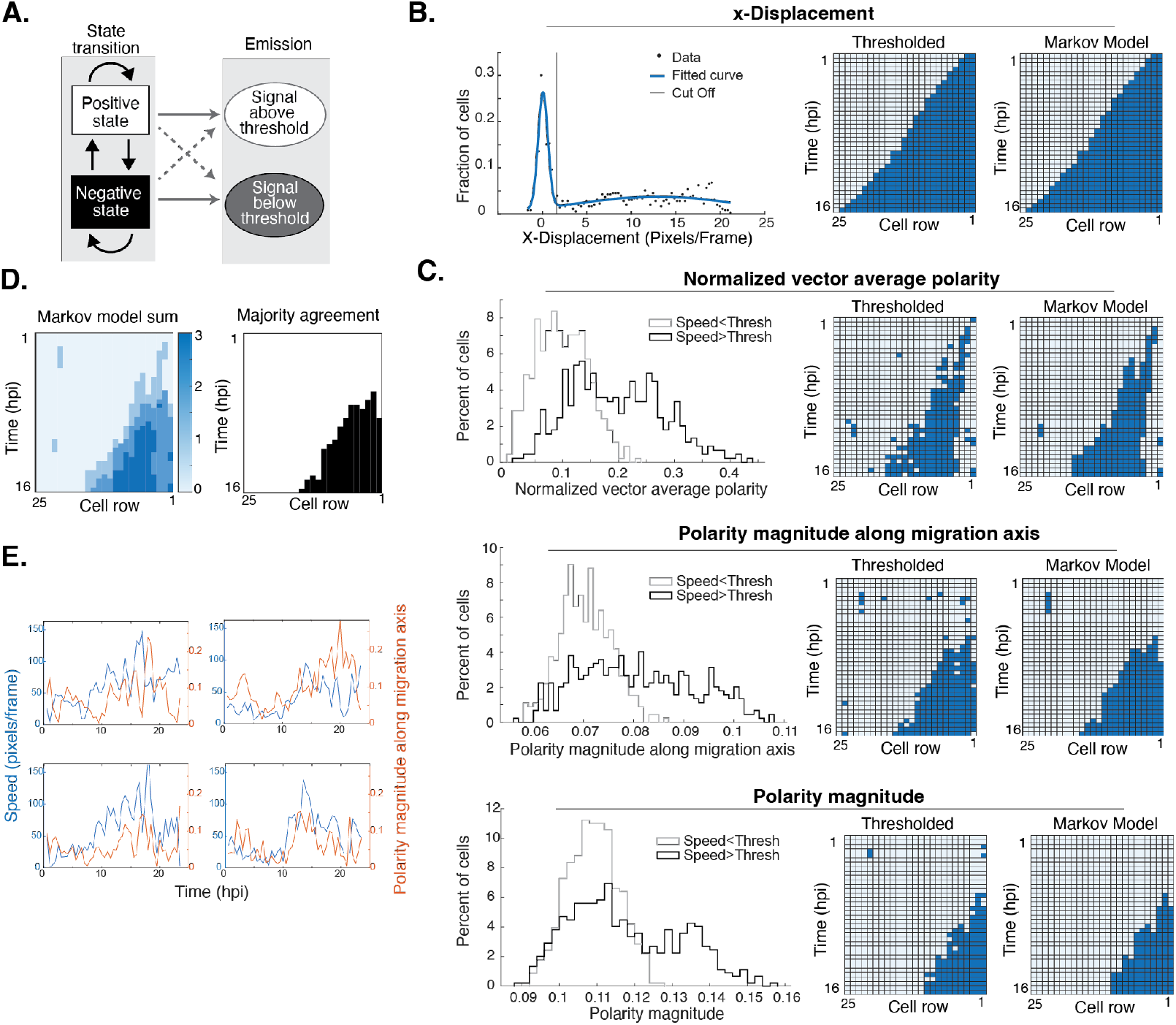
A Markov model for quantifying the timing of migration and polarity initiation. **A**. Depiction of Markov model architecture. Cells were considered positive (migrating or polarized) or negative (nonmigrating or nonpolarized), resulting in bias towards emission above or below determined thresholds. **B**. Determination of the speed threshold above which cells are considered migrating. *Left:* Cells were binned into rows and the x-displacement values of all rows were fit to a two-Gaussian distribution. A value of two standard deviations above the mean of the slower population was used as the cut-off for calling cells migrating vs nonmigrating. *Right*: Kymographs of x-displacement binarized using determined threshold and corresponding Markov model output. **C**. Determination of the polarity thresholds above which an emission is considered polarized. *Left:* Cell rows were binarized as migrating vs nonmigrating based on the speed threshold determined in **B**, and the polarity measurements were compared between the migrating vs nonmigrating rows, including polarity magnitude, polarity magnitude along the migration axis and normalized vector average polarity. As expected, polarity hallmarks are largely private to migrating rows, allowing for use of nonmigrating rows as a true negative when evaluating polarity signal. The polarity measurements among nonmigratory rows were fit to a Gaussian and a polarity threshold was drawn 1.5 standard deviations above the mean. These thresholds were used to binarize data and train a hidden Markov model using the MATLAB Statistics and Machine Learning Toolbox. *Right*: Kymographs of polarity statistics binarized using determined threshold and corresponding Markov model output. **D**. Classification of polarity states. *Left*: Overlay of all polarity Markov model results. Consensus agreement between all three of the described models most consistently described observed polarity; *Right*: Kymograph depicting final polarity determination where rows are called as polarized *(black squares)* if two out of the three polarity metrics are above the respective thresholds. **E**. Representative single cell traces showing both migration speeds and polarity magnitude along the migration axis.

**Figure S5.**
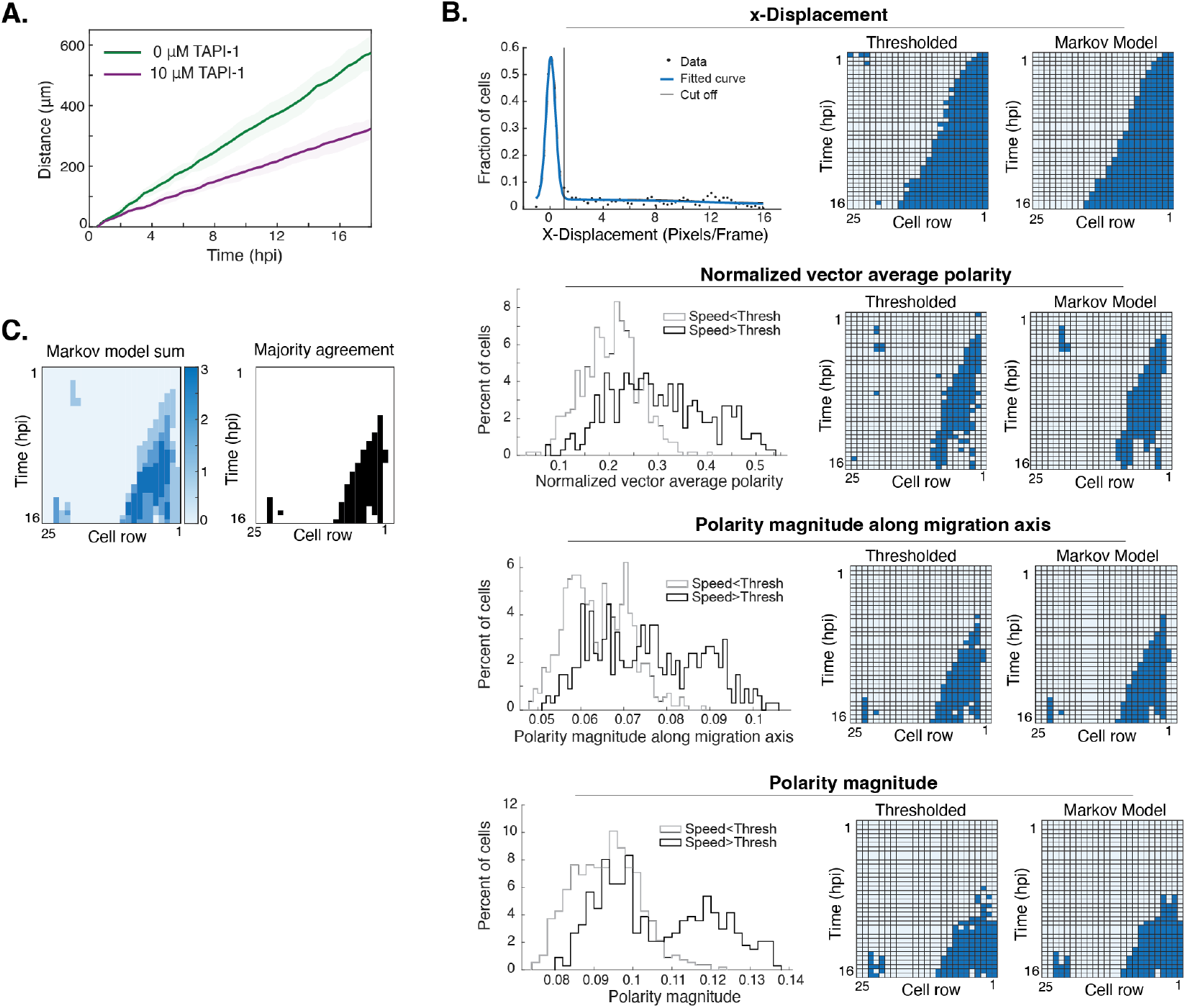
Quantifying the timing of migration and polarity initiation under TAPI-1 treatment. **A**. Speed of migration in untreated and 10 μM TAPI-1 treated cells, as inferred by the displacement of the leading front over time. Lines represent mean and shading indicates SD, 3 replicates. **B-C**. The same thresholding method as in Figure S4B-D was used to binarize data and train the same hidden markov model to determine the migration and polarity states of each cell row, under 10 μM TAPI-1 treatment.

**Figure S6.**
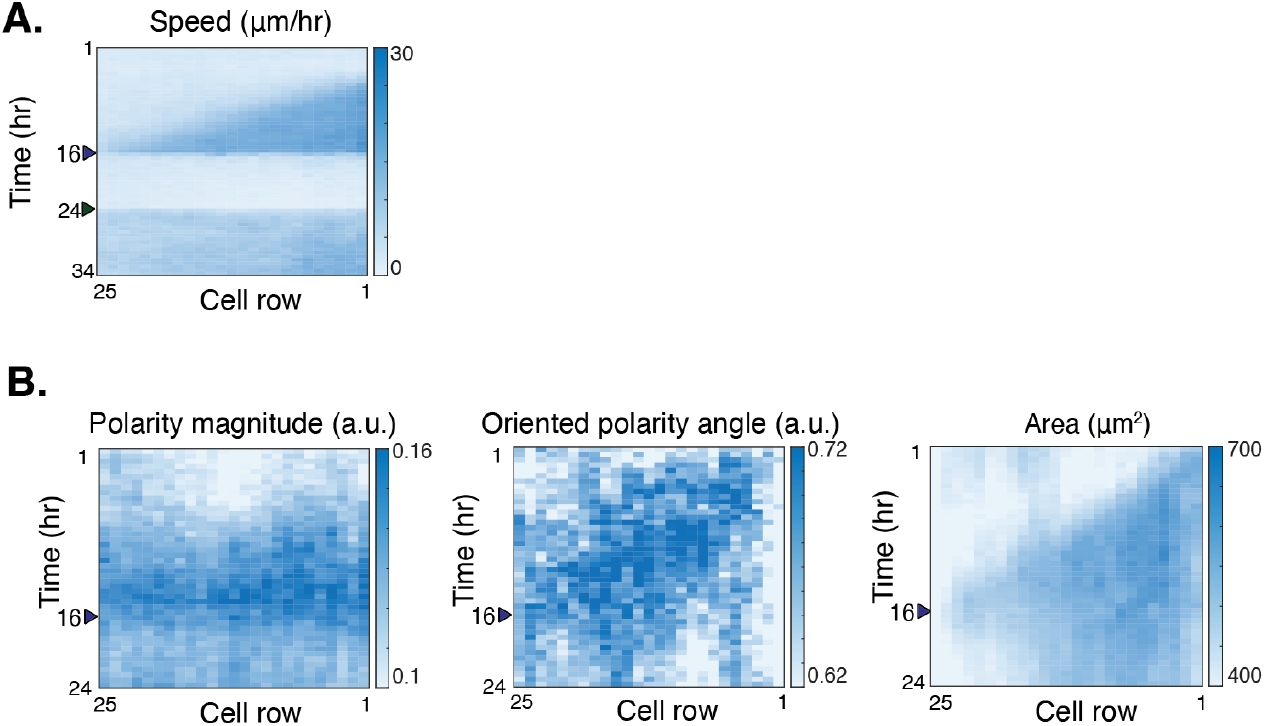
Migration arrest with the ARP2/3 inhibitor is reversible and leads to loss of polarity. **A**. Cell speed before, during and following wash off of CK-666 treatment (5077 cell traces, n=1). **B**. Kymographs of polarity magnitude, oriented polarity angle, and cell area before and during CK-666 treatment (21246 cell traces, n=4). Both polarity magnitude and polarity angle alignment decrease after migration arrest.

**Figure S7.**
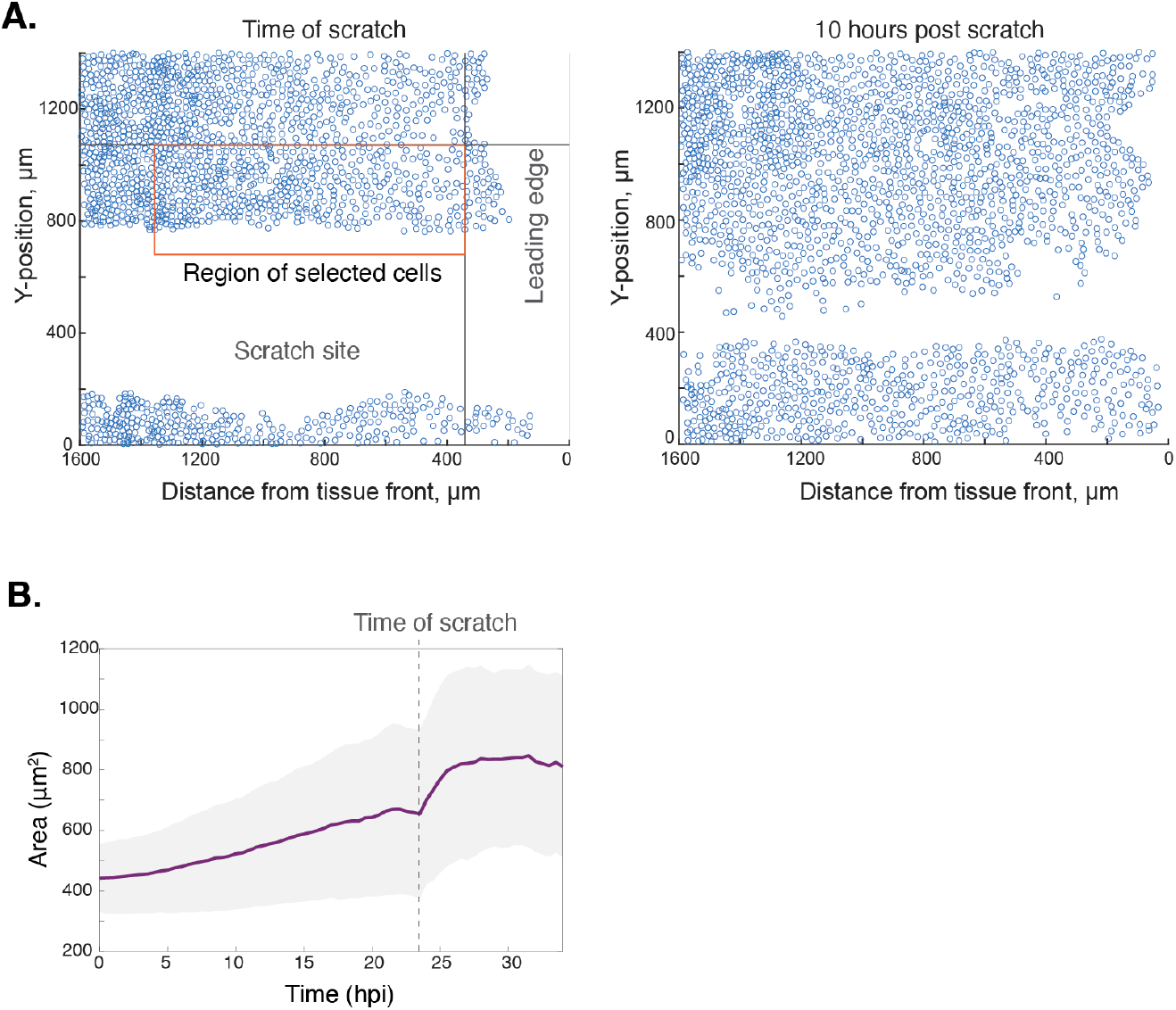
Scratching re-directs cell migration and increases cell area. **A**. Visualization of cell location at time of scratch and 10 hours post scratch. Vertical line indicates the spatial threshold to exclude pre-scratch leading rows (Position > 100 μm from average tissue front), which do not consistently polarize during migration. The horizontal line indicates the spatial threshold to exclude non-reorienting tissue (Position < 300 μm from scratch edge), which was not reached by the new migration wave by the end of the experiment. Cells were additionally filtered on prescratch polarization (magnitude along migration axis > 0.1) and x-displacement using thresholds described as in Fig S4. **B**. Area of selected reorienting cells. Line shows mean, and shading indicates SD (459 cell traces).

**Figure S8.**
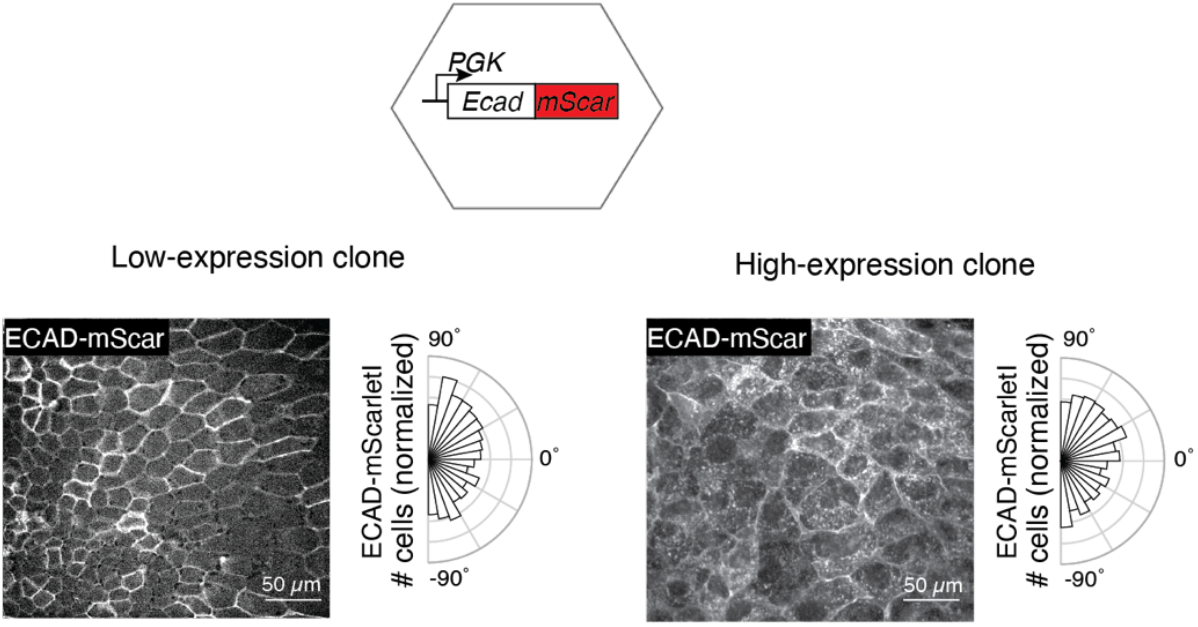
Overexpressed E-cadherin did not polarize during migration. Clonal MDCK cell lines with both low levels *(left)* and high levels *(right)* of E-cadherin-mScarletI overexpression showed E-cadherin junctional localization but failed to polarize (low expression line: 1495 cells, 3 replicates; high expression: 3138 cells, 2 replicates). Cells with higher E-cadherin levels had more internalized mScarletI signal. Representative images of migrating cells at 18 hpi.

**Figure S9.**
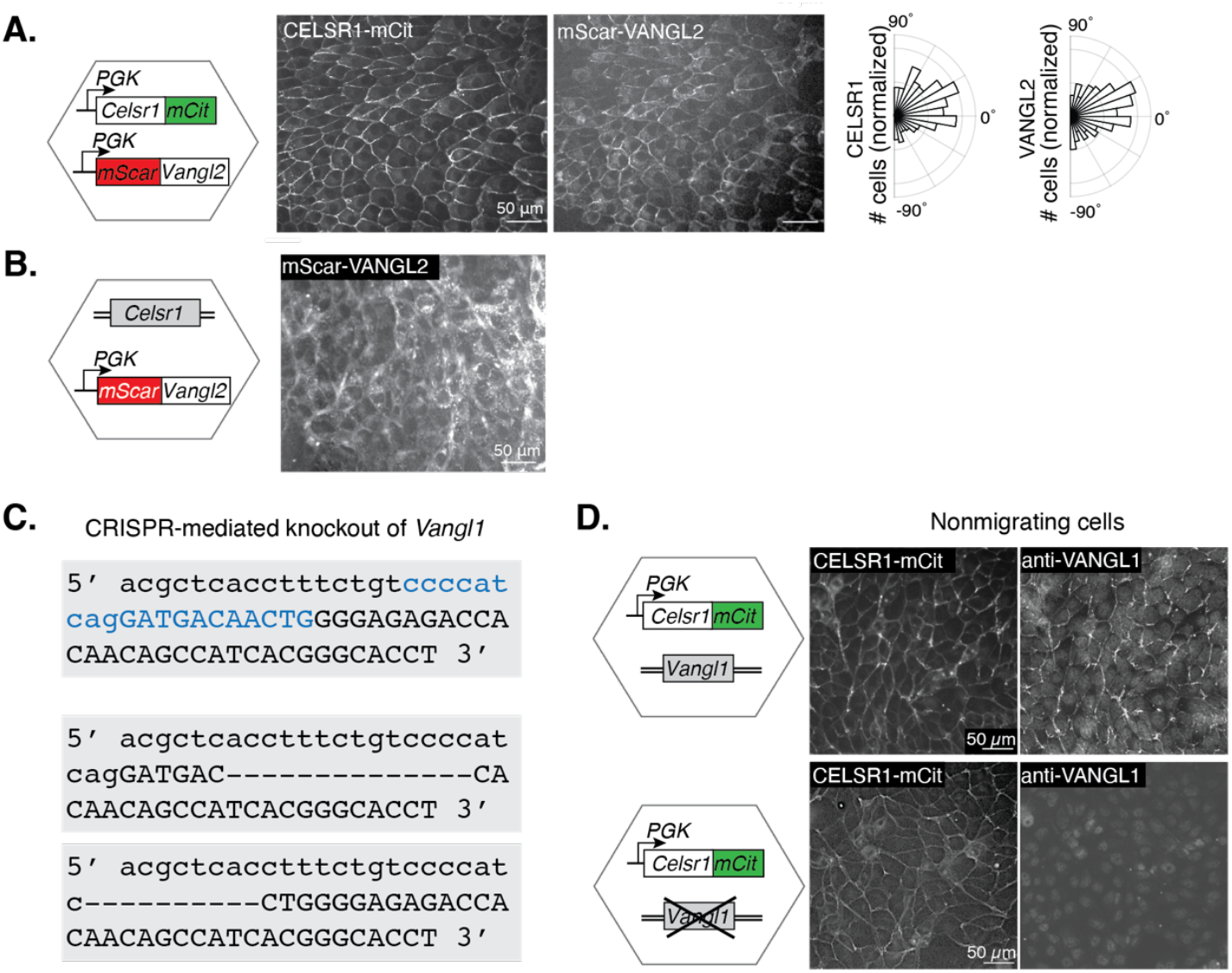
Testing the VANGL-CELSR interaction. **A**. CELSR1-mCitrine/mScarletI-VANGL2 coexpression led to polarization of VANGL2 during migration. Representative images of a migrating field at 18 hpi. **B**. mScarletI-VANGL2 localization in MDCK cells without CELSR1 overexpression. Representative image of a migrating field at 18 hpi. mScarletI-VANGL2 was insufficiently junctionally localized for segmentation and polarity quantification. **C**. CRISPR-mediated knockout of *Vangl1* in CELSR1-mCitrine MDCK cells. Lowercase letters, intron sequence; uppercase letters, exon sequence; blue text, guide RNA target sequence. Wild-type sequence is shown at the top. Allele 1 *(middle)* is a 14-base deletion from the exon, causing a frame shift. Allele 2 *(bottom)* is a 10-base deletion which removes the splice acceptor site. **D**. Staining with anti-VANGL1 antibody showed a loss of VANGL1 protein in *Vangl1* knockout cells. Cells were confluently cultured without induced migration.

**Table S1.**
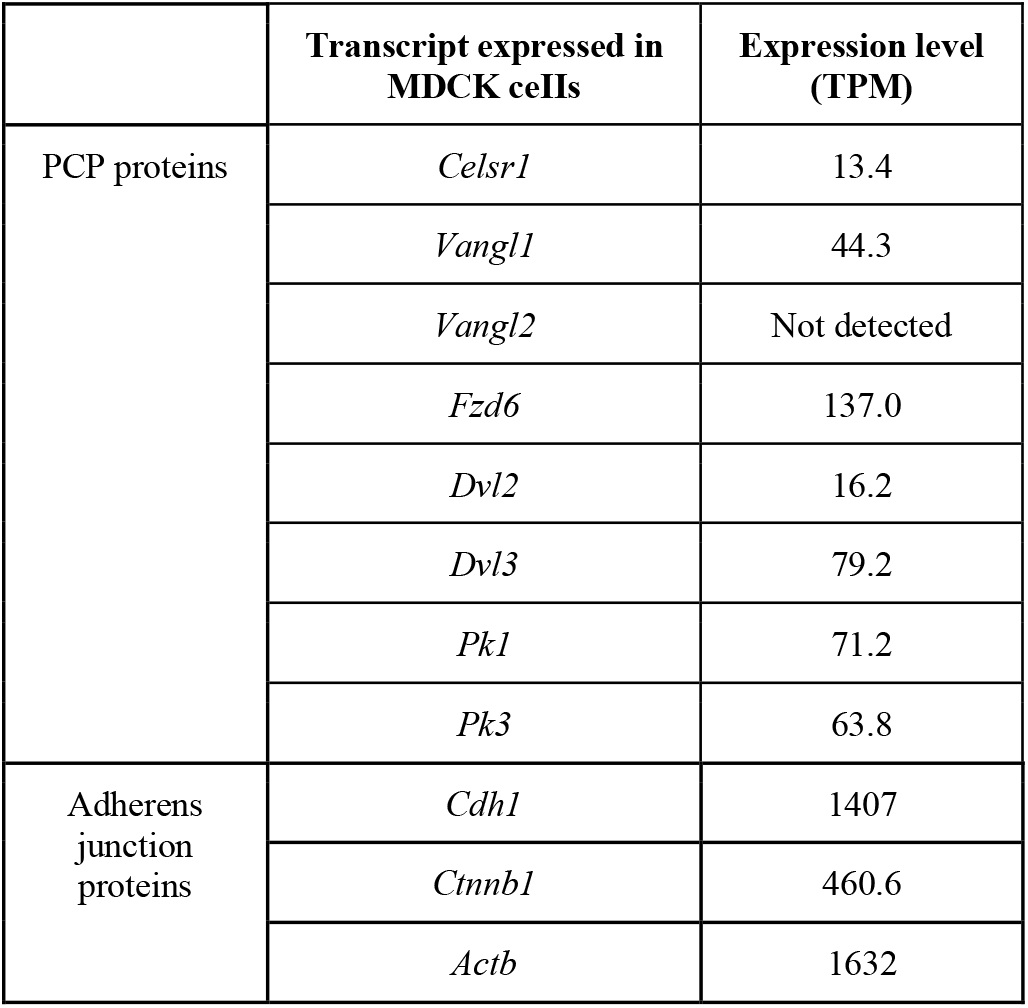
Expression of PCP and other junctional proteins in MDCKs, by bulk RNAseq.

**Table S2.** Binarization thresholds determined from stationary cells.

| **Condition** | **X-displacement threshold ($\mu\text{m/hr}$)** | **Speed threshold ($\mu\text{m/hr}$)** | **Normalized vector average polarity threshold (AU)** | **Magnitude along migration axis threshold (AU)** | **Polarity magnitude threshold (AU)** |
| --- | --- | --- | --- | --- | --- |
| Untreated | 2.09 | 6.86 | 0.207 | 0.0814 | 0.1241 |
| TAPI-1 | 1.39 | 3.75 | 0.33 | 0.0812 | 0.1122 |

**Table S3.** Initial and final model weights for described Markov models.

| Untreated State Transition rates | Initial Training Parameter | Trained X-displacement | Trained speed | Trained normalized vector average polarity | Trained magnitude along migration axis | Trained polarity magnitude |
| --- | --- | --- | --- | --- | --- | --- |
| Off to On | 0.04 | 0.052 | 0.0976 | 0.03444 | 0.0351 | 0.0208 |
| On to Off | 0.01 | 0 | 0.0271 | 0.0488 | 0.0086 | 0 |
| **Untreated Emission Rates** |  |  |  |  |  |  |
| Off emits On | 0.16 | 0 | 0.0271 | 0.0207 | 0.0178 | 0.0185 |
| On emits Off | 0.33 | 0.0014 | 0 | 0.1061 | 0.0406 | 0.0243 |
| **TAPI-1 Treatment State Transition Rates** |  |  |  |  |  |  |
| Off to On | 0.04 | 0.0283 | 0.0216 | 0.0153 | 0.0309 | 0.031 |
| On to Off | 0.01 | 0 | 0.0028 | 0.0933 | 0.0257 | 0.0048 |
| **TAPI-1 Treatment Emission Rates** |  |  |  |  |  |  |
| Off emits On | 0.16 | 0.0128 | 0.0341 | 0.0081 | 0.0003 | 0.0026 |
| On emits Off | 0.33 | 0.0038 | 0 | 0.1209 | 0.0576 | 0.0531 |

**Supplementary Video 1. CELSR1 subcellular distribution is primarily localized to junctions**. Zeiss Airyscan confocal Z-stack of fixed CELSR1-mCitrine MDCK cells at 18 hpi, stained for nuclei (DAPI, blue) and actin (Phalloidin-549, magenta) and visualizing CELSR1-mCitrine (green). Each frame is one slice of a Z-stack from basal to apical with slice thickness 0.13 µm. Still images and orthogonal views are shown in Figure S2.

**Supplementary Video 2. CELSR1 polarity emerges in collectively migrating cells**. Time-lapse video of migrating CELSR1-mCitrine MDCK cells, starting at time of migration induction. Still images shown in Figure 1E.

**Supplementary Video 3. Individual CELSR1-mCitrine cells polarize in response to migration**. Motion-stabilized time-lapse video of migrating CELSR1-mCitrine MDCK cells. Still images shown in Figure 1F.

**Supplementary Video 4. Migration arrest leads to loss of CELSR1 polarity**. Time-lapse video of migrating CELSR1-mCitrine cells before and after the addition of 400 µM CK666 (Arp2/3 inhibitor). CK666 leads to migration arrest and loss of CELSR1-mCitrine+ polarization. Still images shown in Figure 3C.

**Supplementary Video 5. Migrating cells reorient in response to scratch**. Time-lapse video of migration and polarity re-orientation. Migrating CELSR1-mCitrine cells are scratched with a pipette tip at 24 hpi to create an orthogonal leading edge, resulting in migration direction change. Analysis shown in Figure 3D-F.

**Supplementary Video 6. Polarity reorients in individual cells in response to migration direction change**. Motion-stabilized time-lapse video of migrating CELSR1-mCitrine cells after pipette tip scratch. Video starts at 0 hrs post scratch (hps) and follows cells migrating in the newly established orthogonal migration direction. Still images shown in Figure 3G.

## Notes

### Competing Interest Statement

The authors have declared no competing interest.

